# Investigation of ACE2 N-terminal fragments binding to SARS-CoV-2 Spike RBD

**DOI:** 10.1101/2020.03.19.999318

**Authors:** G. Zhang, S. Pomplun, A. R. Loftis, X. Tan, A. Loas, B. L. Pentelute

## Abstract

Coronavirus disease 19 (COVID-19) is an emerging global health crisis. With over 7 million confirmed cases to date, this pandemic continues to expand, spurring research to discover vaccines and therapies. SARS-CoV-2 is the novel coronavirus responsible for this disease. It initiates entry into human cells by binding to angiotensin-converting enzyme 2 (ACE2) via the receptor binding domain (RBD) of its spike protein (S). Disrupting the SARS-CoV-2-RBD binding to ACE2 with designer drugs has the potential to inhibit the virus from entering human cells, presenting a new modality for therapeutic intervention. Peptide-based binders are an attractive solution to inhibit the RBD-ACE2 interaction by adequately covering the extended protein contact interface. Using molecular dynamics simulations based on the recently solved cryo-EM structure of ACE2 in complex with SARS-CoV-2-RBD, we observed that the ACE2 peptidase domain (PD) α1 helix is important for binding SARS-CoV-2-RBD. Using automated fast-flow peptide synthesis, we chemically synthesized a 23-mer peptide fragment of the ACE2 PD α1 helix (SBP1) composed entirely of proteinogenic amino acids. Chemical synthesis of SBP1 was complete in 1.5 hours, and after work up and isolation >20 milligrams of pure material was obtained. Bio-layer interferometry (BLI) revealed that SBP1 associates with micromolar affinity to insect-derived SARS-CoV-2-RBD protein obtained from Sino Biological. Association of SBP1 was not observed to an appreciable extent to HEK cell-expressed SARS-CoV-2-RBD proteins and insect-derived variants acquired from other vendors. Moreover, competitive BLI assays showed SBP1 does not outcompete ACE2 binding to Sino Biological insect-derived SARS-CoV-2-RBD. Further investigations are ongoing to gain insight into the molecular and structural determinants of the variable binding behavior to different SARS-CoV-2-RBD protein variants.

## 1. Introduction

A novel coronavirus (SARS-CoV-2) from Wuhan, China, has caused over 7 million confirmed cases and over 400,000 deaths globally, according to the COVID-19 situation report from WHO on June 10, 2020 (https://www.who.int/emergencies/diseases/novel-coronavirus572019/situation-reports/), and the number is continually growing. Similar to the SARS-CoV outbreak in 2002, SARS-CoV-2 causes severe respiratory problems. Coughing, fever, difficulties in breathing and/or shortage of breath are the common symptoms. Aged patients with pre-existing medical conditions are at most risk with a mortality rate ~1.5% or even higher in some regions. Moreover, human-to-human transmission can occur rapidly by close contact. To slow this pandemic and treat infected patients, rapid development of specific antiviral drugs is of the highest urgency.

The closely-related SARS-CoV coronavirus invades host cells by binding the angiotensin65 converting enzyme 2 (ACE2) receptor on human cell surface through its viral spike protein (S) [1–4]. It was recently established that SARS-CoV-2 uses the same receptor for host cell entry and binds ACE2 with an affinity comparable with the corresponding spike protein of SARS-CoV [5, 6]. Recent cryo-electron microscopy (cryo-EM) structural studies of the SARS-CoV-2 spike protein receptor binding domain (RBD) in complex with full-length human ACE2 receptor revealed key amino acid residues at the contact interface between the two proteins and estimated the binding affinity at ~15 nM [7, 8]. These studies provide valuable information that can be leveraged for the development of disruptors specific for the SARS-CoV-2/ACE2 protein-protein interaction (PPI). Small-molecule inhibitors are often less effective at disrupting extended protein binding interfaces [9]. Peptides, on the other hand, offer a synthetically accessible solution to disrupt PPIs by binding at interface regions containing multiple contact “hot spots” [10].

We hypothesized that disruption of the viral SARS-CoV-2-RBD/host ACE2 interaction with peptide-based binders would prevent virus entry into human cells, offering a novel opportunity for therapeutic intervention. To investigate this hypothesis, we launched a campaign to design and discover minimum-length peptide binders to SARS-CoV-2-RBD. Notably, coronavirus spike proteins have been previously targeted with peptide-based fragments, such as an extended ACE2 helical mimic developed against SARS-CoV-RBD [11], pan-CoV inhibitors of the fusion of the spike S2 subunit with the cell membrane [12, 13], and an 85-mer N-terminal truncate of the ACE2 protein that binds SARS-CoV-2-RBD with nanomolar affinity [14]. Complementing these approaches, our efforts aimed to determine the minimum length required for an ACE2 N-terminal peptide fragment that maintains specific association with SARS-CoV-2-RBD.

Analyzing the cryo-EM structure of the SARS-CoV-2-RBD/ACE2 complex, we found that the binding interface spans a large elongated surface area, as is common for PPIs. We leveraged molecular dynamics simulations and automated fast-flow peptide synthesis [15] to prepare a 23-mer peptide binder (SBP1) to SARS-CoV-2-RBD, the sequence of which was derived from the ACE2 α1 helix. Using bio-layer interferometry (BLI), we determined that N-terminal biotinylated SBP1 binds Sino Biological insect-derived SARS-CoV-2-RBD with micromolar affinity (dissociation constant, *K*_D_ = 1.3 μM). We also found, however, that N-terminal biotinylated SBP1 does not associate with HEK-expressed SARS-CoV-2-RBD or insect-derived variants purchased from other commercial sources. Although biotinylated SBP1 binds Sino Biological insect-derived SARS-CoV-2-RBD when immobilized on BLI streptavidin tips, no specific disruption of the SARS-CoV-2-RBD/ACE2 interaction was observed in solution in a BLI competition assay. Further investigation is currently ongoing to understand the different association behaviors of this ACE2-derived peptide to SARS-CoV-2-RBD protein variants.

## 2. Results

### Molecular dynamic simulations guide peptide binder design

Using the Amber force field[16] a helical peptide sequence (spike-binding peptide 1, SBP1) derived from the α1 helix of ACE2 peptidase domain (ACE2-PD) in complex with SARS-CoV-2-RBD was simulated under TIP3P explicit water conditions. Analyzing the simulation trajectory after 200 ns, we found that SBP1 remains on the spike RBD protein surface in a stable conformation (Fig. 2B) with overall residue fluctuations smaller than 0.8 nm compared with their starting coordinates (Fig. 2A). Per-residue analysis along the 200 ns trajectory showed that the middle residues of SBP1, a 12-mer sequence we termed SBP2, have significantly reduced fluctuations (Fig. 2C, 2D), indicating key interactions. The results of this MD simulation suggest that SBP1 and SBP2 peptides derived from the ACE-PD α1 helix may alone potentially bind the SARS-CoV-2 spike RBD protein with sufficient affinity to disrupt the associated PPI.

**Figure 1.**
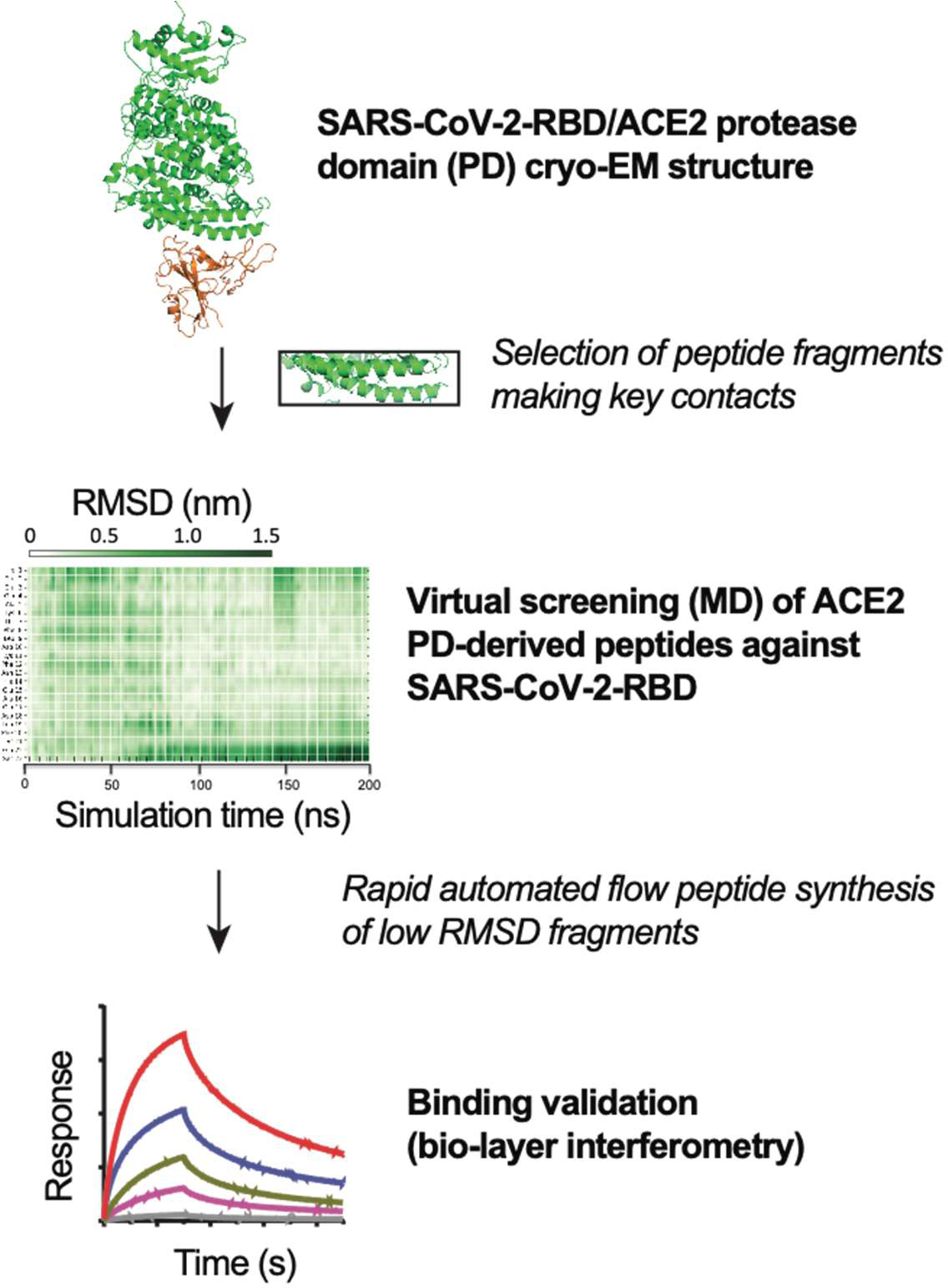
MD-guided target selection for rapid flow synthesis of a SARS-CoV-2-RBD297 binding peptide. Fragments of ACE2-PD domain are docked against SARS-CoV-2 receptor-binding domain (PDB: 6M17). Low RMSD peptides are rapidly synthesized by fully automated flow peptide synthesis, and binding to glycosylated SARS-CoV-2-RBD is determined by BLI.

**Figure 2.**
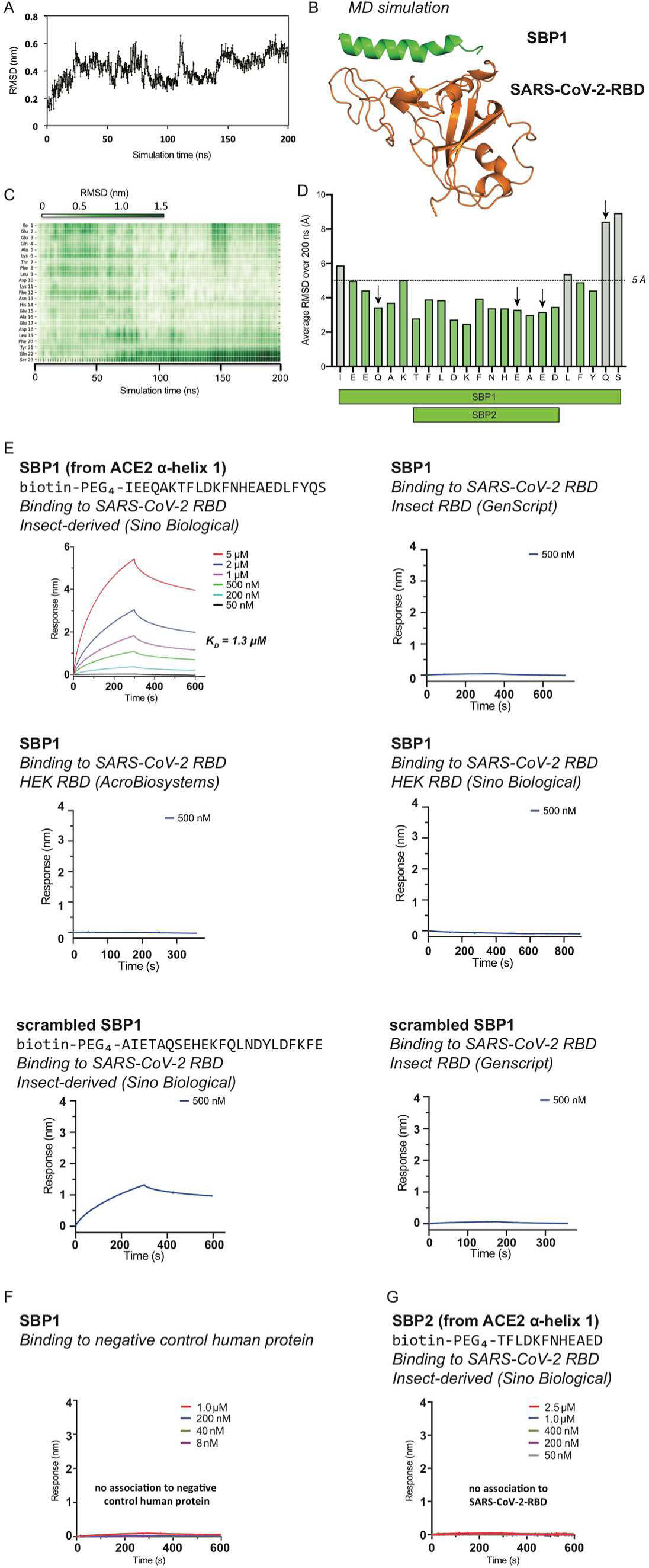
Human ACE2-PD domain α-helix 1-derived SBP1 binds only Sino Biological insect-derived SARS-CoV-2-RBD. (A) RMSD for SBP1 docked to SARS-CoV-2-RBD during 200 ns MD simulation. (B) Binding interface between SARS-CoV-2-RBD and SBP1 after 200 ns simulation. Individual RMSD (C) and average RMSD (D) values for SBP1 residues over the course of the 200 ns simulation. Arrows indicate residues contributing key hydrogen bonding interactions (determined using UCSF Chimera, Version 1.12). Individual residues with RMSD below 5 Å arbitrarily colored green. (E) Binding affinity of SBP1 and a scrambled SBP1 sequence to various sources of glycosylated SARS-CoV-2-RBD proteins determined by bio-layer interferometry (BLI). (F) BLI association response of SBP1 to negative control human protein menin and (G) BLI association response of 12-mer SBP2 peptide to Sino Biological insect-derived SARS-CoV-2-RBD.

### Automated fast-flow peptide synthesis yields >95% pure compound

The two N-terminal biotinylated peptides, SBP1 and SBP2, derived from the α1 helix were prepared by automated fast-flow peptide synthesis[15, 17] with a total synthesis time of 1.5 h over 35 coupling cycles. After cleavage from resin, global deprotection, and subsequent C18 solid115 phase extraction, the purity of the crude peptides was estimated to be >95% for both biotinylated SBP1 and SBP2 based on LC-MS TIC chromatograms (Supplemental Fig. 1). We assessed this purity as acceptable for direct downstream biological characterization.

### SBP1 peptide binds Sino Biological insect-derived SARS-CoV-2-RBD with micromolar affinity, but does not associate with other commercial sources of SARS-CoV-2-RBD

Bio-layer interferometry (BLI) was used to measure the binding affinity of the synthesized peptide SBP1 to glycosylated Sino Biological insect-derived SARS-CoV-2-RBD, Sino Biological HEK-expressed SARS-CoV-2-RBD, GenScript insect-derived SARS-CoV-2-RBD and AcroBiosystems HEK-expressed SARS-CoV-2-RBD. In all of these assays, biotinylated SBP1 was immobilized onto streptavidin (SA) biosensors. After fitting the association and dissociation curves from serial dilutions of the protein, the dissociation constant (*K*_D_) of SBP1 to glycosylated Sino Biological insect-derived SARS-CoV-2-RBD was determined to be ~1300 nM using the global fitting algorithm and 1:1 binding model (Fig. 2E). However, SBP1 did not associate with the other three SARS-CoV-2-RBD proteins studied (Fig. 2E). Surprisingly, a scrambled sequence of SBP1 exhibited binding to the Sino Biological insect-derived SARS-CoV-2-RBD with comparable association response to SBP1 at 500 nM concentration (Fig. 2E). SBP1 had no observable binding to a negative control human protein menin (Fig. 2F). Likewise, no association was observed between the biotinylated 12-mer SBP2 and Sino Biological insect-derived SARS-CoV133 2-RBD (Fig. 2F).

### SBP1 does not compete with biotinylated ACE2 binding to Sino Biological insect-derived SARS-CoV-2-RBD

Using a competition-format BLI assay, we confirmed that soluble human ACE2 protein could compete with immobilized biotinylated ACE2 (AcroBiosystems) for binding Sino Biological insect-derived SARS-CoV-2-RBD, and that a 5-fold excess of soluble ACE2 (relative to immobilized biotinylated ACE2) abolished nearly all of the initial ACE2/RBD binding interaction (Fig. 3B, 3D). However, competition was not observed when using non-biotinylated SBP1 pre141 mixed in solution with Sino Biological insect-derived SARS-CoV-2-RBD, even with a 1000-fold excess of the peptide (Fig. 3C, 3E). These data suggest that SBP1 potentially binds SARS-CoV143 2-RBD at a different site than ACE2, binds SARS-CoV-2-RBD too weakly, or for other unknown reasons cannot disrupt the native ACE2/RBD interaction.

**Figure 3.**
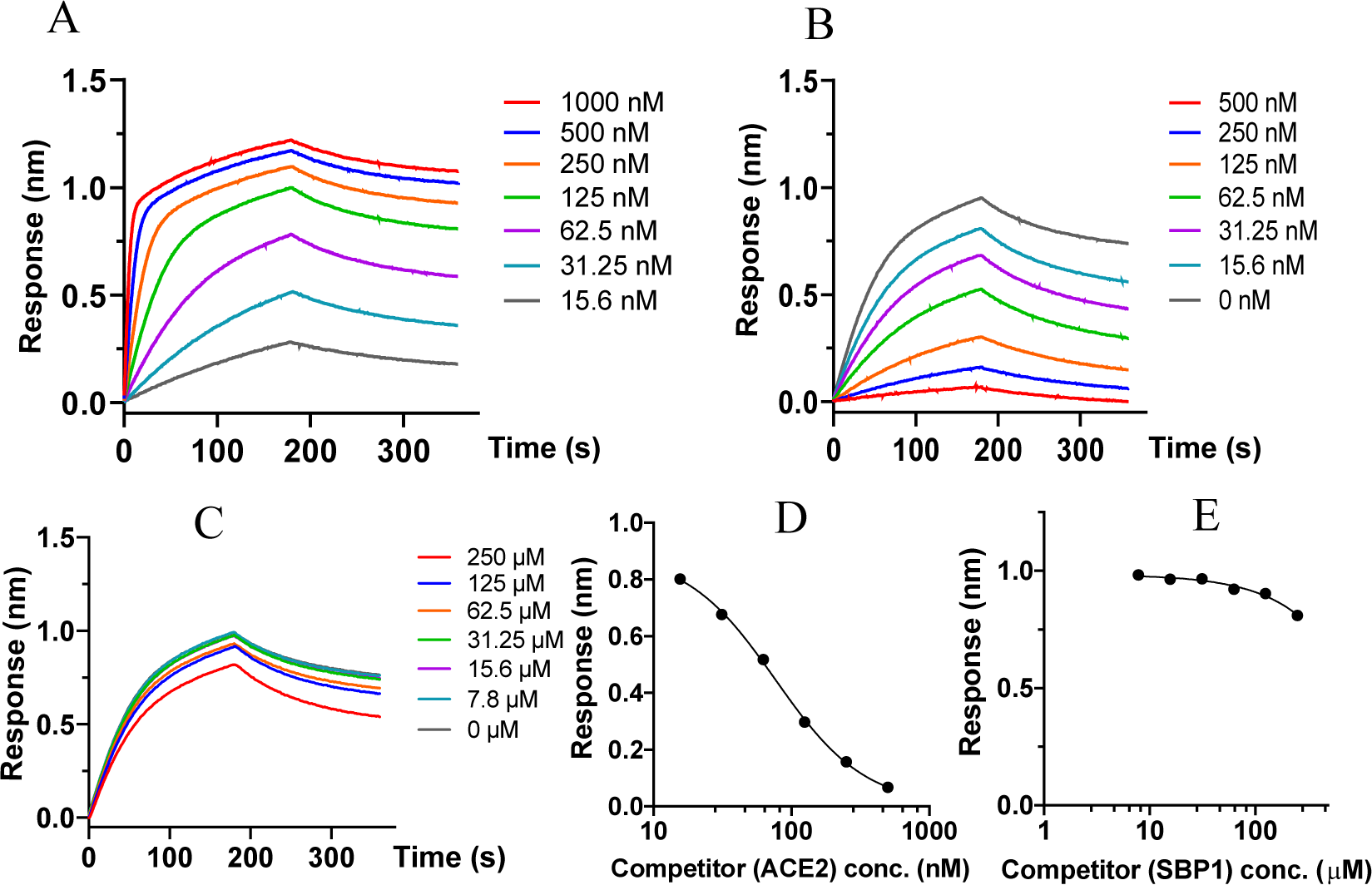
SBP1 does not compete with ACE2 binding to Sino Biological insect-derived SARS-CoV-2-RBD. (A) Association responses of biotinylated ACE2 to Sino Biological insect derived SARS-CoV-2-RBD at different concentrations. The kinetic dissociation constant determined under these conditions was *K*_D_, _ACE2_ = 15 nM. (B and D) Association responses of biotinylated ACE2 with Sino Biological insect-derived SARS-CoV-2-RBD (kept constant at 100 nM) after mixing with soluble human ACE2 at different concentrations. (B) shows the BLI traces and (D) shows re-plots of the endpoint association response (nm) as a function of ACE2 concentration. (C and E) Association responses of biotinylated ACE2 with Sino Biological insect-derived SARS-CoV-2-RBD (kept constant at 100 nM) after mixing with SBP1 at different concentrations. (C) shows the BLI traces and (E) shows re-plots of the endpoint association response (nm) as a function of SBP1 concentration.

## 3. Discussion

Recently published cryo-EM structures of the RBD of SARS-CoV-2 in complex with human ACE2 have identified this PPI as a key step for the entry of SARS-CoV-2 into human cells [7, 8]. Blocking this binding interface represents a highly promising therapeutic strategy, as it could potentially hinder cellular uptake of SARS-CoV-2 and intracellular replication.

Drugging PPIs is a longstanding challenge in traditional drug discovery and peptide-based approaches might help to solve this problem. Small molecule compounds are unlikely to bind large protein surfaces that do not have distinct binding pockets. Peptides, on the other hand, display a larger surface area and chemical functionalities that can mimic and disrupt the native PPI, as is the case for the clinically approved HIV peptide drug Fuzeon [18, 19].

The identification of a suitable starting point for drug discovery campaigns can be time156 intensive. During a pandemic such as this one, therapeutic interventions are urgently needed. Peptide-based strategies were developed to target both the spike protein RBD and S2 subunit of the first SARS-CoV virus [11,12]. Translating these approaches to SARS-CoV-2, inhibitors of the spike protein fusion with the cell membrane and engineered mini-proteins that bind SARS-CoV-2 RBD were developed [13,14]. We aimed to determine the minimum length required of the ACE2 N-terminal peptide fragment in order to maintain binding affinity to SARS-CoV-2-RBD and thus potentially deliver a synthetically accessible therapeutic candidate. To rapidly identify potential short peptide binders to the SARS-CoV-2 spike protein, we used molecular dynamics (MD) simulation on peptides extracted from the human ACE2 sequence. The starting point of the binding simulations was the cryo-EM model of the SARS-CoV-2 spike protein and several peptides derived from the SARS-CoV-2-spike binding domain of human ACE2 protein. Our MD simulation (200 ns trajectory) indicated that the SBP1 peptide, corresponding to the N-terminal ACE2 α1 helix, stably bound to SARS-CoV-2-RBD. The overall peptide fluctuations were smaller than 0.8 nm from the starting coordinates (Fig. 2A). These results indicated the potential of identifying a short SARS-CoV-2-RBD-binding peptide derived from the human ACE2 α1 helix.

A 23-mer peptide sequence (SBP1) was synthesized by automated flow peptide synthesis [15]. The 23 residues selected from the ACE2 α1 helix sequence (IEEQAKTFLDKFNHEAEDLFYQS) showed low fluctuations along the MD simulation trajectory and several important interactions with the spike protein were observed consistently with multiple lines of published data [8, 20]. We used this peptide (SBP1) as an experimental starting point for the development of a SARS-CoV-2 spike protein binder. Our rapid automated flow peptide synthesizer enabled the synthesis of tens of milligrams of SBP1 peptide within 1.5 h. The crude purity was determined to be >95% and therefore sufficient for binding validation by BLI.

The interaction between N-terminal biotinylated SBP1 and the RBD of glycosylated SARS180 CoV-2 spike protein was investigated in detail. We performed serial dilutions of the soluble protein to reliably determine the binding affinity of SBP1 to Sino Biological insect-derived SARS-CoV-2-RBD. Using a global fitting algorithm, we found that N-terminal biotinylated SBP1 binds Sino Biological insect-derived SARS-CoV-2-RBD with micromolar affinity (*K*_D_ = 1.3 <M), a value almost 100-times higher than the estimated binding affinity of the native ACE2 receptor (*K*_D_ ~ 15 nM [7]) (Fig. 2E). The decreased binding affinity relative to ACE2 may partially explain why SBP1 was unable to significantly disrupt ACE2 binding to Sino Biological insect-derived SARS-CoV-2-RBD even at 1000-fold excess in a BLI competition assay (Fig. 3C,E). In addition, the comparable affinity observed for a scrambled sequence of SBP1 (Fig. 2E) suggests the possibility of a promiscuous peptide binding site on the surface of glycosylated Sino Biological insect-derived SARS-CoV-2-RBD at a different location than the one involved in ACE2 receptor binding.

An important outcome of our preliminary BLI binding studies relates to the significantly different SBP1 association behavior observed toward the various SARS-CoV-2-RBD commercial sources investigated. Among the four variants (two insect-derived and two HEK-expressed), Sino Biological insect-derived SARS-CoV-2-RBD was the only one displaying observable association with SBP1 in the BLI Octet assay. This behavior may be partially ascribed to variability in the site196 specific patterns and surface density of SARS-CoV-2-RBD post-translational modifications (PTMs), in particular glycosylation [21], imparted by the different biological expression sources. Distinct PTM patterns are evident in the different mass distribution envelopes obtained by deconvolution of the Sino Biological insect-derived and HEK-expressed SARS-CoV-2-RBD protein total ion current bands, respectively, analyzed by LC-MS (Supplemental Fig. 2). Access to non-glycosylated SARS-CoV-2-RBD either by bacterial recombinant expression or total chemical synthesis in flow [15] may provide additional insight into the role of glycans in modulating peptide interactions with the SARS-CoV-2-RBD/ACE2 interface. Investigations along these lines are in progress to gain additional understanding of these molecular processes.

In conclusion, a biotinylated peptide sequence derived from human ACE2 was found to bind Sino Biological insect-derived SARS-CoV-2 spike protein RBD with micromolar affinity, but did not associate with SARS-CoV-2-RBD variants obtained from other commercial sources. In spite of this association, competitive BLI data indicates that SBP1, even at 1000-fold excess, did not compete with ACE2 for binding to SARS-CoV-2-RBD. Our preliminary studies highlight the unexpected challenges researchers may encounter while developing peptide-based approaches to disrupt the specific interactions of SARS-CoV-2 with its mammalian cell membrane receptors. At the same time, the BLI experiments draw attention to the wide variability in the behavior of the SARS-CoV-2 spike protein variants in solution, likely a consequence of the biological expression source, manufacturing and/or formulation protocols. The development of peptide-based disruptors of SARS-CoV-2 cell entry effective in a clinical setting relies on a broader evaluation and understanding of its spike protein isoforms.

## 4. Experimental Materials and Methods

### GPU-accelerated molecular dynamic simulation

The cryo-EM structure of ternary complex of SARS-CoV-2-RBD with ACE2-B^0^ 219 AT1 (PDB: 6M17) was chosen as the initial structure, which was explicitly solvated in an 87 Å^3^ box, to perform a 200 ns molecular dynamical (MD) simulation using NAMD on MIT’s supercomputing clusters (GPU node). The Amber force field was used to model the protein and peptide. The MD simulation system was equilibrated at 300 K for 2 ns. Periodic boundary conditions were used and long-range electrostatic interactions were calculated with particle mesh Ewald method, with non bonded cutoff set to 12.0 Å. SHAKE algorithm was used to constrain bonds involving hydrogen atoms. Time step is 2 fs and the trajectories were recorded every 10 ps. After simulation production runs, trajectory files were loaded into the VMD software for further analysis.

### Automated fast-flow peptide synthesis

SBP1 and SBP2 sequences were synthesized at 90 °C on Rink Amide-ChemMatrix resin with HATU activation using a fully automatic flow-based peptide synthesizer. Amide bond formation was performed in 8 seconds, and Fmoc groups were removed in 8 seconds with 40% (v/v) piperidine in DMF. The overall synthesis cycle was completed in ~120 seconds per amino acid incorporated. After completion of fast-flow synthesis, the resins were washed with DMF (3 x) and then incubated with HATU-activated biotin-PEG_4_-propionic acid (CAS# 721431-18-1) at room temperature for 1.0 h for biotinylation on the peptide N-terminus.

### Peptide cleavage and deprotection

After peptide synthesis, the peptidyl resin was rinsed with dichloromethane briefly and then dried in a vacuum chamber overnight. Next day, approximately 5 mL of cleavage solution (94% trifluoroacetic acid (TFA), 1% TIPS, 2.5% EDT, 2.5% water) was added into the syringe containing the resin. The syringe was kept at room temperature for 2 h before injecting the cleavage solution into a 50 mL conical tube. Dry-ice cold diethyl ether (~50 mL) was added to the cleavage mixture and the precipitate was collected by centrifugation and triturated twice with cold diethyl ether (50 mL). The supernatant was discarded. Residual ether was allowed to evaporate and the peptide was dissolved in water with 0.1% TFA for solid-phase extraction.

### Solid-phase extraction (SPE)

After peptide cleavage, peptide precipitates were dissolved in water with 0.1% TFA. Agilent Mega BE C18 column (Part No: 12256130) was conditioned with 5 mL of 100% acetonitrile with 0.1% TFA, and then equilibrated with 15 mL of water with 0.1% TFA. Peptides were loaded onto the column for binding, followed by washing with 15 mL of water with 0.1% TFA, and finally, eluted with 5 mL of 30/70 water/acetonitrile (v/v) with 0.1% TFA.

### Liquid chromatography-mass spectrometry (LC-MS)

Peptides were dissolved in water with 0.1% TFA followed by LC-MS analysis on an Agilent 6550 iFunnel ESI-Q-ToF instrument using an Agilent Jupiter C4 reverse-phase column (2.1 mm × 150 mm, 5 µm particle size). Mobile phases were 0.1% formic acid in water (solvent A) and 0.1% formic acid in acetonitrile (solvent B). Linear gradients of 1 to 61% solvent B over 15 minutes (flow rate: 0.5 mL/min) were used to acquire LC-MS chromatograms.

### Kinetic binding assay using bio-layer interferometry (BLI)

A ForteBio Octet^®^ RED96 Bio-Layer Interferometry system (Octet RED96, ForteBio, CA) was used to characterize the in vitro peptide-protein binding affinity at 30 °C and 1000 rpm. Briefly, streptavidin (SA) tips were dipped in 200 μL of biotinylated peptide solution (2.5 μM in 1x kinetic buffer: 1x PBS with 0.1% BSA and 0.05% tween) for the loading step. The tips loaded with peptide were then sampled with four different sourced SARS-CoV-2-RBD proteins (Sino Biological insect263 derived RBD, CAT: 40592-V08B; Sino Biological HEK-expressed RBD, CAT: 40592-V08H; AcroBiosystems HEK-expressed RBD, CAT: SPD-C52H3; GenScript insect-derived RBD, CAT: Z03479) or menin protein at various concentrations in 1x kinetic buffer to obtain the association curve. Peptide only was used as reference for background subtraction. After association, the tips were dipped back into 1x kinetic buffer to obtain the dissociation curve. The association and dissociation curves were fitted with ForteBio Biosystems using four experimental conditions (n = 6, global fitting algorithm, binding model 1:1) to obtain the dissociation constants *K*_D_.

### In-solution BLI competition assay

A BLI competition binding assay was set up as previously described.[22] First, a calibration curve was constructed using biotinylated ACE2 dipping into a serially-diluted Sino Biological insect275 derived SARS-CoV-2-RBD. In brief, streptavidin sensors were soaked in kinetic buffer (PBS supplemented with 0.02% Tween-20, and 0.1% BSA) for 10 min at 30 °C, and loaded with biotinylated ACE2 for 4 min. Then, serial dilutions of SARS-CoV-2-RBD in Kinetic buffer were analyzed for binding, typically at 30 °C and 1,000 rpm. Second, with the binding information from the calibration curve, a concentration of SARS-CoV-2-RBD protein at 100 nM was chosen to premix with various concentration of SBP1 to study the competition effects. Briefly, different concentration of SBP1 was incubated with 100 nM Sino Biological insect-derived SARS-CoV-2-RBD protein at room temperature for 15 min. Meanwhile, streptavidin sensors were soaked into kinetic buffer for 10 min at 30 °C. ACE2 was immobilized on the streptavidin sensor surface and the association and dissociation curves of SARS-CoV-2-RBD in the preincubated samples were then analyzed at 30 °C and 1,000 rpm.

## Acknowledgements

The authors thank the MIT Supercomputing Center for providing the computational resources to run MD simulations. This research was supported by a COVID-19 Fast Grant award sponsored by Emergent Ventures at the Mercatus Center, George Mason University, and by MIT seed funds. S.P. is supported by a postdoctoral fellowship from Deutsche Forschungsgemeinschaft (DFG, award PO 2413/1-1). MIT has filed a provisional patent application related to this work.

## Competing interests

B.L.P. is a founder of Resolute Bio and Amide Technologies.

---

**Supplemental Figure 1.**
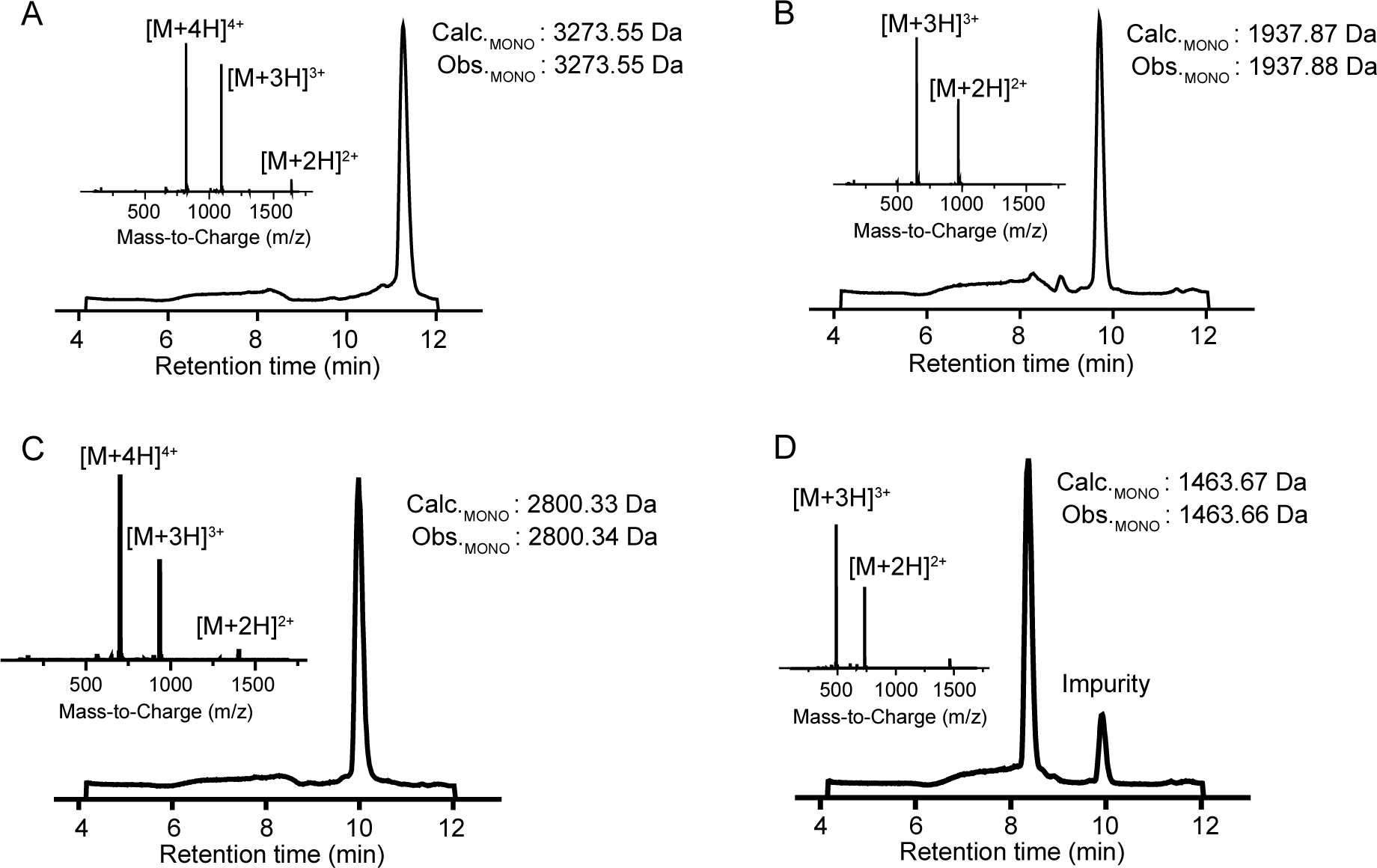
ACE2-derived peptides were prepared by solid-phase peptide synthesis. Total ion current chromatograms (TIC) and associated mass spectra of purified N terminal biotinylated SBP1 peptide (A), purified N-terminal biotinylated SBP2 peptide (B), crude SBP1 peptide (C), and crude SBP2 peptide (D).

**Supplemental Figure 2.**
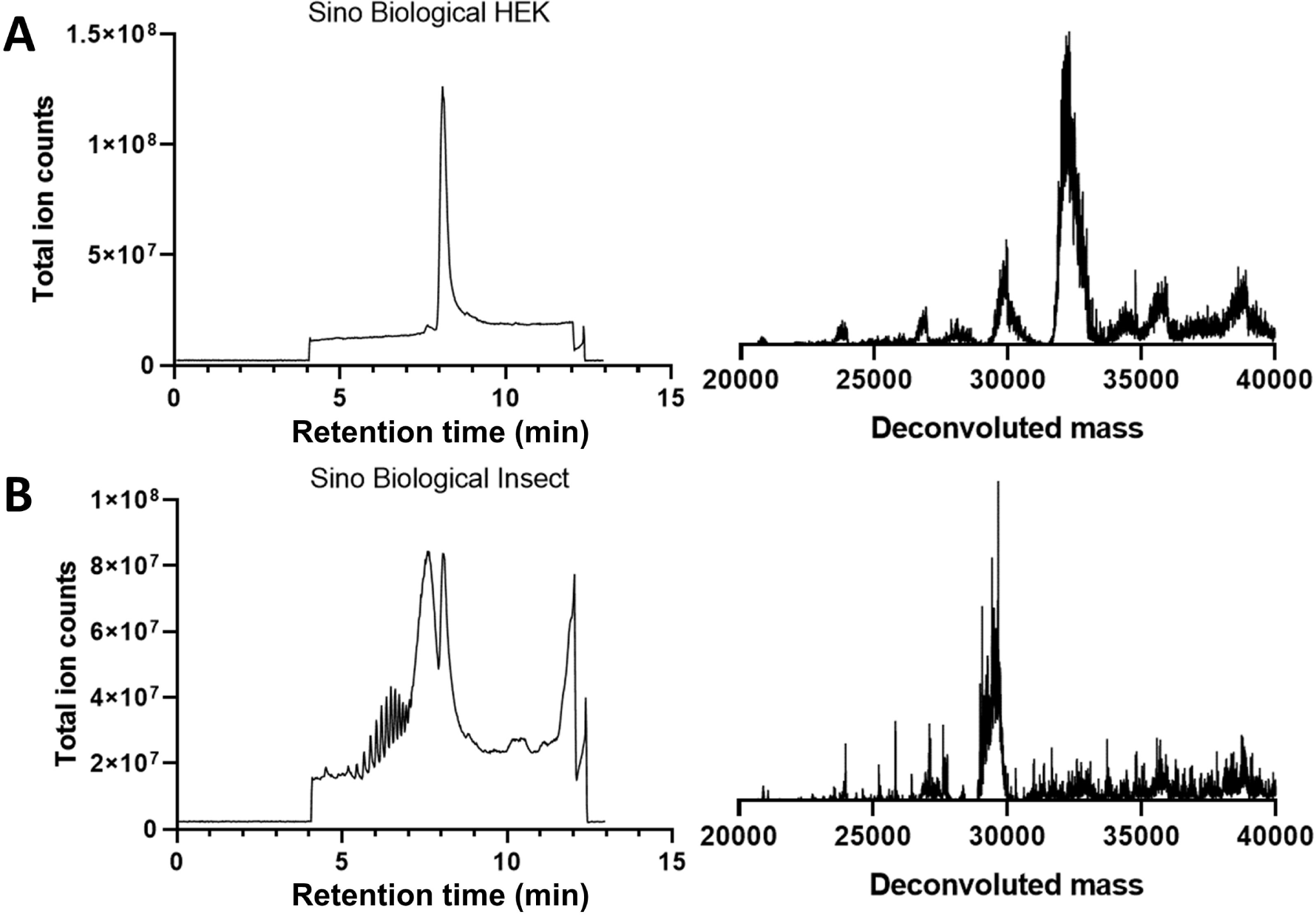
SARS-CoV-2 spike protein RBD domains derived from different biological sources display differences in mass distribution profiles. Total ion current chromatograms (TIC) obtained by LC-MS analysis of commercial samples of (A) glycosylated Sino Biological HEK-expressed SARS-CoV-2-RBD (solution in phosphate-buffered saline) and (B) glycosylated Sino Biological insect-derived SARS-CoV-2-RBD with associated deconvoluted mass spectra obtained by integration over the protein peak at ~8 min. The broad bands in the TIC chromatograms of (B) are due to additives present in the vendor-formulated solid powder (10% glycerol, 5% trehalose, 5% mannitol and 0.01% tween-80).

## Notes

### Competing Interest Statement

Bradley L. Pentelute is a co-founder of Resolute Bio and Amide Technologies.

### Summary of Updates

We revised and expanded our binding affinity studies of our lead SBP1 peptide with various SARS-CoV-2-RBD isoforms obtained from different commercial sources and added the following experiments. Using bio-layer interferometry (BLI), we determined that N-terminal biotinylated SBP1 binds Sino Biological insect-derived SARS-CoV-2-RBD with micromolar affinity. We also found, however, that N-terminal biotinylated SBP1 does not associate with HEK-expressed SARS-CoV-2-RBD or insect-derived variants purchased from other commercial sources. Although biotinylated SBP1 binds Sino Biological insect-derived SARS-CoV-2-RBD when immobilized on BLI streptavidin tips, no specific disruption of the SARS-CoV-2-RBD/ACE2 interaction was observed in solution in a BLI competition assay. Title, abstract and author list was also updated to reflect the new studies and findings.

